# Evolutionary biomechanics of maximum running speed in spiders (Araneae)

**DOI:** 10.64898/2026.06.11.731532

**Authors:** Shreyas Kuchibhotla, Michael Kelly, Vincent Jackel, Elizabeth Bane, Hendrik K. Beck, Jonas O. Wolff, David Labonte

## Abstract

**Background:** Maximum running speed is a central performance trait, linking morphology, physiology and behaviour to fitness. It is shaped by physical capacity and ecological selection but may also be constrained by ancestry. To examine how these forces interact across macroevolutionary timescales, we conducted an allometric study in a hyper-diverse arthropod taxon—spiders (Araneae).

**Results:** Drawing on running performance data for 258 species from 68 of the 139 extant spider families, we integrated phylogenetic comparative methods and biomechanical modelling to disentangle the effects of body size, ancestry, leg morphology, ecological guild and preferred locomotor orientation. Maximum running speed varied substantially, both across body mass and among species of similar body mass. By accounting for body mass with a biomechanical model, we show that size-specific performance carries a strong phylogenetic signal, and that high-performing runners first evolved within the derived infraorder Araneomorphae. Strong running performance, after accounting for both body size and shared ancestry, was associated with relatively longer legs and, to a lesser extent, ecological guild, but not with leg slenderness or a preference for inverted versus upright locomotion.

**Conclusions:** Macroevolutionary patterns of running performance thus reflect not only variation in body size, but also size-specific leg morphology, ecological differentiation and phylogenetic history. We hope this study contributes to the development of formal evolutionary biomechanics—one that seeks to explain patterns of diversity through the explicit integration of large-scale comparative data, natural history and quantitative models derived from first principles.

## Introduction

Morphology, physiology and behaviour can influence fitness through what they enable organisms to do—run quickly, bite strongly, move efficiently, or cling securely. Whole-organism performance can therefore be seen as a bridge between form and fitness [1–3]. Selection, however, does not act on an unconstrained “design” space. Instead, phenotypic evolution is channelled by phylogenetic history, developmental pathways, and physical laws, and, across macroevolutionary time, diversification in performance is thus the result of interactions among multiple, and sometimes competing, selective pressures and constraints [4–6]. Disentangling this complexity requires broad comparative data and reliable measures of maximum performance, but great care is required where performance is expected to co-vary with body size, because absolute performance can obscure relevant variation or even be misleading altogether -it is arguably not particularly informative to observe that an elephant can lift more load than an ant in absolute terms. Instead, it is the *size-specific* performance that can reveal species that are unusually fast, slow, strong or weak, and that thus carries biologically, and not just physically, meaningful information [7–11]. But how should one account for body size?

It is here that biomechanics, no doubt a sub-discipline in its own right, becomes an indispensable component of macroevolutionary analyses [12]. The key strength of biomechanical analyses is that they can complement statistical models with predictions grounded in first principles [7, 8, 13]. In some cases, this can make size-correction straightforward. The maximum force an animal can generate, for example, is proportional to the maximum isometric stress and the physiological cross-sectional area of its muscles. Stress is typically assumed to be size-invariant, and the area of muscle, under the parsimonious assumption of geometric similarity, is proportional to *m*^2*/*3^, where *m* is body mass. The maximum force, then, should scale as *m*^2*/*3^, too, a classic prediction that has robust empirical support [14]. Thus, if an animal ten times as heavy produces about 10^2*/*3^ *≈* 5 times as much force, one is likely looking at an effect primarily of size; but if the difference is much smaller or larger, other factors may be at play.

Unfortunately, things are not always so simple. A notorious counterexample is maximum running speed, a trait linked to escape performance, prey capture success, dispersal, territoriality and reproductive competition, and thus of substantial ecological relevance [15–18]. Unlike many performance traits, running performance does not appear to follow a simple monotonic power law [18, 19]. Instead, the fastest animals are of intermediate size, a pattern that continues to attract biomechanical debate [18, 20–23], and that renders physically-grounded size correction challenging. Empirically, there is little room for doubt that the top speed of a cheetah exceeds that of a hippopotamus. But does this difference arise primarily from size-dependent physical limits, from phylogenetic history, from ecological differences in selective pressures, or from some weighted interaction among these factors?

Recently, we have put forward a biomechanical model that predicted the unusual allometry of maximum running speed observed across Animalia both qualitatively and quantitatively [21, 24]. By combining the fundamental principle of the conservation of energy with constraints imposed by muscle physiology, we showed that muscle has not one, but two characteristic energy capacities [24, 25]. Because these two capacities scale differently with size, small animals tend to be limited by one, whereas large animals are more likely limited by the other -the resulting allometry of running speed becomes non-trivial, and cannot be described by a classic allometric power law with just a single exponent [21, 24]. A biomechanical model that captures this complexity, rests on simple first-principles arguments, and enjoys strong empirical support, creates a new opportunity to assess the extent to which variation in running speed is shaped by phylogenetic history, morphology, and ecology.

To implement such an approach, we focus on the Araneae, a clade that combines broad phylogenetic diversity with striking ecological and locomotor variation. Species differ in whether they build burrows, tube webs, substrate webs or aerial webs; whether they hunt on the ground, in foliage or even underwater; and whether they move predominantly upright (on substrates) or inverted (on silk threads) [26–28]. These differences may well alter both the selective value of running speed, and the way in which it is realised. Spiders also combine muscle-driven leg flexion with hydraulically assisted leg extension [29, 30]. We therefore ask whether a muscle-based model can account for running-speed variation despite this mixed actuation mechanism. Against this backdrop, we ask four questions:

1. Does maximum running speed in spiders follow a simple allometric relationship, or are the fastest spiders of intermediate size [21]?
2. How much of the variation in size-corrected running speed can be explained by variation in relative leg length and leg slenderness [26]?
3. After accounting for body size and phylogenetic history, are differences in running speed related to ecological niche [31]?
4. Does size-corrected running speed vary systematically across the spider tree of life?

To address these questions, we combine comparative phylogenetic methods with biomechanical modelling and analyse maximum running speed in a sample comprising 258 species from 68 of the 139 extant spider families, selected to capture the taxonomic and ecological diversity of extant spiders, and with particular effort to include some of the smallest and largest taxa within major evolutionary lineages.

## Materials and methods

### Study animals

We measured maximum sustained running speed for 246 specimens representing 162 species. To maximise phylogenetic coverage, we supplemented these measurements with published data for a further 96 species [31–45], yielding a final dataset of 258 species from 68 of the 139 extant spider families, covering about six orders of magnitude in body mass. A complete species list is provided in the supplementary information.

Most specimens measured in this study were collected in and around London and Greifswald from October 2023 to November 2025 (permit number VG-23-055, where necessary). A further 30 species were sourced through fieldwork in North America, Southern Europe and Australia (permit numbers SF/0114/24 and 476/2024/CAPT, where necessary), and 24 species were purchased from, or recorded directly at, pet suppliers. Species were chosen to represent a sufficiently broad taxonomic diversity and size range. We also aimed to record species from a variety of ecological groups to avoid bias towards any specific hunting strategies. Among the individuals contributing maximum-speed records for the newly measured species, 102 were adults, of which 39 were males. Adult male spiders have a different morphology from juveniles and adult females and may often be more adapted for mate search [40, 46, 47]. To test for a potential bias, we repeated the main analyses after exclusion of adult males; the qualitative conclusions were unchanged, and we hence pooled the data across sexes and life stages, to maximize taxon sampling across our models. Owing to limited availability, some species were represented by one specimen only (see supplementary information).

Although there are no formal guidelines on non-cephalopod invertebrates in the 1986 Animals (Scientific Procedures) Act [48], measures were taken to ensure minimal harm during recordings, as well as adequate husbandry, in accordance with the Royal Entomological Society’s guidelines [49]. Specimens were kept in ventilated boxes under controlled conditions, and with species-appropriate substrates. UK-collected specimens were maintained at 25*^◦^*C, whereas specimens housed in Greifswald were kept at 20– 25*^◦^*C for tropical and thermophilic species, 20–22*^◦^*C for temperate species, and 6*^◦^*C for cavernicolous species. Animals were fed small arthropods from laboratory colonies, and provided with water by regular misting, via damp cotton balls, or small dishes.

Specimens were usually euthanised soon after experiments by freezing; some were preserved after natural death or by directly placing them in 70–100% ethanol. Both methods are widely adopted for euthanising and preserving invertebrates [50, 51].

### Running speed assays

Maximum sustained running speed was defined as the highest mean speed maintained for more than a few isolated strides. Running speed assays for all but three species were conducted on horizontal A4 or A3 grid paper, which was mounted on a rigid plastic tray or metal sheet to provide both a high-traction substrate and a scale reference; liquid paraffin, thinly applied to the walls, prevented escape by climbing species. All trials were filmed with a camera positioned perpendicular to the running plane (Fig. 1), with illumination provided by two LED lights or desk lamps placed along the upper edges of the enclosure. Three large theraphosid specimens were recorded instead on an automated laboratory treadmill [52], which allowed multiple strides to be filmed with a single camera. Trial temperature was typically between 21-23*^◦^*C, but never dropped below 18 or exceeded 26*^◦^*C.

**Fig. 1.**
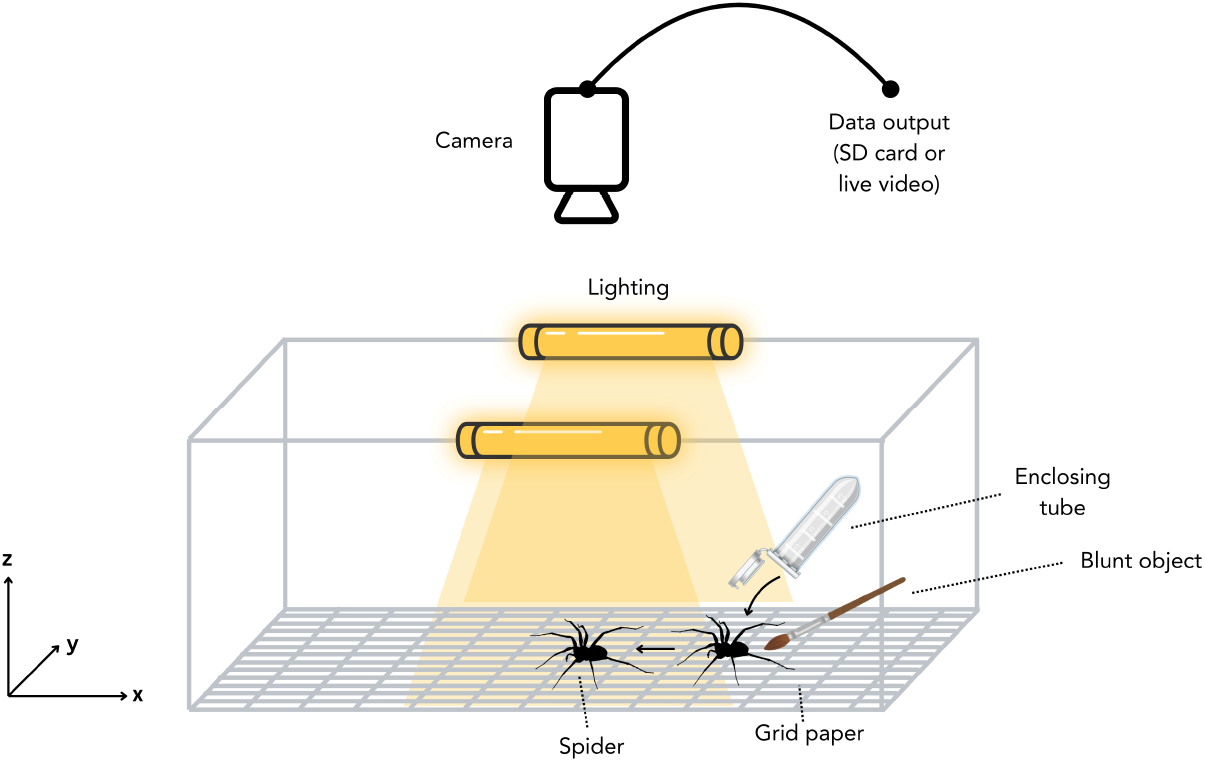
Schematic of the recording setup (not to scale). Spiders were placed on a grid sheet of known dimensions. An escape response was then induced by touching the abdomen with an object such as a paintbrush or the blunt end of a pencil and recorded with a high-speed camera from the dorsal aspect to measure maximum sustained running speed.

Recordings were made using a FLIR Blackfly BFS-U3-51S5C-C equipped with a Kowa LM25JC 25 mm lens, mounted on a custom-built frame, or using tripod-mounted cameras (Sony Cybershot RX10, Nikon D850 with a Sigma 14–24 mm F2.8 DG DN lens, or a Basler Ace CMOS Monochrome with a Fujinon HF12.5HA-1B lens). Frame rates ranged from 60 to 500 fps, and were controlled using SpinView (Spinnaker SDK, FLIR Integrated Imaging Solutions Inc., Richmond, BC, Canada) or TroublePix (Norpix Inc., Montreal, Canada) as appropriate. The distance between the camera and recording plane was adjusted to specimen size, yielding fields of view spanning 0.1-1.6m. The FLIR setup exhibited slight barrel distortion, which was corrected in MATLAB R2024b (MathWorks, Natick, MA, USA) using built-in camera-calibration and pointundistortion routines from the Computer Vision Toolbox (estimateCameraParameters, undistortPoints), and photographs of a 12.5mm checkerboard. Slight camera tilt, identified *post hoc* by visual inspection in seven recordings, was corrected using built-in geometric-transformation routines from the Image Processing Toolbox (fitgeotrans, transformPointsForward), and the known dimensions of the grid squares.

Before each trial, spiders were weighed to *±*0.1 mg (ABJ 120-4MN, Kern & Sohn, Balingen-Frommern, Germany, or Ohaus Explorer, Parsippany, USA) or, for large species, to *±*0.1 g (Kern PCB 10000-1 or kitchen scale); they were then placed in a tube on the recording substrate. After the tube was removed, some spiders ran spontaneously; in most cases, however, an escape response was elicited by gentle contact with a paintbrush, the blunt end of a pencil, or by puffs of compressed air. Because true maximal performance is hard to elicit reliably in the laboratory, escape responses are widely used as a practical proxy for near-maximal performance [16, 53–55]. For each specimen, 3–10 trials were recorded to increase the likelihood of capturing maximal locomotor performance.

Raw videos were inspected for each species individually to identify the fastest trial, typically from the largest individual or, where body mass differed little among specimens, the fastest trial overall. Where the fastest run was uncertain, several candidate trials were taken forward for analysis. In each selected trial, the posterior end of the carapace, a proxy for the body centre of mass, was tracked using either OmniTrax with manual refinement [56]; manual tracking in Blender v3.3; or OpenCV v4.6. This combination of methods was necessitated by the extremely broad range of spider sizes and speeds; any automated tracks were checked before analysis. For some small species, this point was not consistently visible, so the centre of the spider’s silhouette was tracked instead. Acceleration was readily apparent in the tracked frames through the notable increase in distance covered across equal consecutive time intervals, so those intervals were excluded. Sequences of typically 5–50 frames were selected manually after initial acceleration, with approximately constant speed checked visually and by comparing distances covered per frame. The number of frames depended on the animal’s size, speed, and stamina. Maximum sustained running speed was then calculated as the distance travelled during these strides divided by elapsed time, using the 10-mm grid on the A3 paper for spatial calibration. For our measurements, this procedure selected sustained running after initial acceleration and avoided averaging over prolonged bouts in which speed declined.

To extend the taxonomic coverage of our dataset, we included presumed maximum running speeds from the literature [31–45]. We cannot establish beyond reasonable doubt whether every included data point represents sustained speed or the species’ true maximal capacity. Our working assumption is that submaximal measurements mainly add scatter rather than a systematic bias with body size. For the 80 species from Kelly et al. [31], body mass was estimated from body length, BL, as *m*_mg_ = 0.18BL^2.70^, following Penell et al. [regression on 189 European spider species, *R*^2^ = 0.96; see 57].

### Morphometry

Morphometric data comprised leg I and leg IV length (excluding coxa and trochanter), tibia widths at the midpoint, carapace width, and carapace length. Legs I and IV were measured to reduce dependence on a single leg pair. The front and hind legs vary most in dimensions across spiders [28], and the sum of both determines the total effective leg span, a key determinant of stride length. Because tibia and femur width covary strongly in spiders [28], we measured only tibia width at its midpoint, preferred because the tibia more closely approximates a cylinder than the femur, whose width can vary substantially along its length. Leg segments were measured from the approximate endpoint of the proximal joint to the base of the distal joint to *±* 0.01 mm, with their sum rounded to 0.1 mm to account for cumulative measurement error. Carapace width was taken as the greatest distance perpendicular to the anterior–posterior axis, or, in damaged specimens, as twice the distance from the fovea to the carapace margin. Carapace length was measured as the shortest distance between the anterior median eyes and the base of the pedicel; where this point was obscured by the abdomen, the endpoint was estimated from the assumed convergence of the symmetrical carapace margins. From these data, we derived three parameters for further analyses: relative leg length (mean leg length (I+IV)/carapace length); leg slenderness (mean leg length (I+IV)/mean tibia width (I+IV)); and carapace area, estimated by assuming that carapace width and length form the two axes of an ellipse.

All measurements were taken from photographs of live specimens (immobilised or anaesthetised), or preserved specimens (dry or in ethanol). Specimens were photographed in a Petri dish under a Leica Z6 microscope with 0.8*×* and 5*×* objective lenses (Leica Microsystems Ltd, Milton Keynes, United Kingdom), with scale provided either by an AGL4078 UKAS certified stage micrometer, or, for some specimens, by a ruler placed beneath the dish. The carapace was photographed in dorsal view, with legs dissected or repositioned as necessary to obtain a planar orientation. Wherever possible, legs were removed from the body and photographed in lateral view. In some specimens, leg removal was avoided because of the rarity of the material. In these cases, the body was positioned to allow lateral or dorsal photographs of the legs, which were gently extended and held in place using tweezers, gel, or damp tissue. For three theraphosids, it was not possible to obtain leg dimensions on the live animals, so instead, exuviae were photographed on 10 mm grid paper with the carapace in dorsal view and detached legs in lateral view. To correct for the size change upon moulting, live spiders were photographed dorsally with a ruler, and exuvia-based leg measurements were then scaled by the ratio of live to moult carapace length, assuming geometric similarity. Similarly, two large araneomorphs were anaesthetised with CO_2_ and photographed on grid paper. Enabled by bilateral symmetry [58], measurements were taken from limbs of one side only, chosen at random. All measurements were completed in ImageJ v1.53 [59].

Two subadult males that moulted before morphometric measurements could be completed were excluded from further analysis. For 47 species, the specimens used in running assays were not available for morphometric measurements, as spiders were kept alive for other experiments, or specimens were damaged; the dataset was thus supplemented with measurements from the literature [60–79], the World Spider Trait Database [80], or from similarly-shaped conspecifics instead. Where even this was not possible (*Philodromus praedatus*, *Oonops pulcher*, *Pardosa nigriceps*, *Xenesthis immanis*, *Xenesthis/Pamphobeteus sp. “megascopula”*, *Kibramoa sp.*, *Selenops sp. “Arizona”* and *Phrurolithus festivus*), we used published measurements for similarly-sized congeners (*Philodromus aureolus*, *Oonops gavarrensis*, *Pardosa lugubris*, *Xenesthis intermedia*, *Kibramoa suprenans*, *Selenops submaculatus* and *Phrurolithus flavipes*) [80–83]. After phylogenetic matching, full carapace and leg measurements were available for 123 assayed specimens; the supplemented dataset covered 176 species, including 157 with leg widths. For scaling analysis on carapace area, data where mass was estimated from body length—related to carapace size—were removed (45 species).

### Phylogenetic inference and guild categorisation

To enable phylogenetic comparative analyses, a phylogeny was inferred from molecular data via maximum likelihood in IQ-TREE v2 [84]. To construct the molecular matrix, sequences of sections of three mitochondrial genes (12S, 16S, COI) and three nuclear genes (18S, 28S, H3) -called ‘markers’ in the following -were obtained from the NCBI GenBank or the Barcode of Life Database [85], the literature [31, 86–88], or, for 28 species, assembled *de novo* using the protocol by Gajski et al. [89]. For each gene, data were assembled and edited in Geneious [90] and MEGA v11 [91], and subsequently aligned using ClustalW. Marker sequences with outstanding length were trimmed, and obviously misaligned or disparate sequences were checked for contamination via BLAST searches against the NCBI database and removed as necessary. Maximum likelihood gene trees were inferred using default settings in the W-IQ-TREE web application [92].

Our sample spans the entire spider tree of life, but a small number of marker genes are unsuitable to resolve deep time nodes. To obtain a stable phylogenetic backbone, we included contigs with thousands of ultraconserved elements (UCE) for a subsample of 120 species from our taxon sample or close relatives [31, 87, 93, 94]; 25 additional species without trait data were added to enrich the phylogenetic backbone, or to serve as outgroup lineages to root the tree. The UCE matrix was created from assembled contigs from the studies by Kulkarni et al. [94], Kelly et al. [31, 93], and Wolff et al. [87] via phyluce [95], using 25% gene occupancy with strict GBlocks cleaning (b1 = 0.5, b2 = 0.85, b3 = 4, b4 = 8).

For phylogenetic inference, each UCE locus and each marker was treated as a separate partition. Molecular model selection, maximum-likelihood phylogenetic inference, and bootstrapping were performed in IQ-TREE 2 [84], using the HPC Brain Cluster of the University of Greifswald, with branch support assessed via ultra-fast bootstrap (ufb) [96] and SH-aLRT branch tests [97]. The resulting tree was converted to an ultrametric chronogram using the chronos function in the R package ape [98]. Secondary node calibrations were taken from the minimum and maximum ages of the 95% highest posterior density intervals for 24 dating points in the fossil-calibrated spider chronogram of [99], mapped to the corresponding tree nodes; the root age followed [28]. Species lacking trait data, included originally to enrich the backbone, were dropped from the phylogeny after the time calibration. The final trait dataset contained 258 species, of which 243 could be matched to terminal taxa in the pruned chronogram used for phylogenetic comparative analyses. The pruned ultrametric tree used for these analyses is provided in the supplementary information.

To supplement phylogenetic information with ecological context, species were assigned one of six guilds and one of two dominant modes of locomotion (upright or inverted), using prior coding for the species or the closest available related taxon [28, 31, 80, 100–108]. Guild 1, burrow/tube web builders, comprises spiders living in burrows or tube-shaped webs in the ground or other substrates; guild 2, substrate web builders, comprises spiders building webs directly onto a substrate; guild 3, aerial web builders, comprises spiders building two-or three-dimensional webs primarily suspended in the air; guild 4, ground runners, comprises cursorial spiders actively hunting on the ground; guild 5, foliage hunters, comprises cursorial spiders actively hunting on plants or similar vertical substrates; and guild 6, ambush hunters, comprises non-web-building spiders waiting for prey rather than pursuing it. For further details on guild classification, see [28]. Semi-aquatic taxa such as Zoicinae and Dolomedidae were classified as ground runners, and the two web-invading araneophagic spiders—*Ero aphana* and *Ero furcata* (Mimetidae)—were classed as guild 3, since they behave and locomote like web-building spiders [109]. Upright and inverted refer to the predominant mode of locomotion in nature, and species from both locomotor categories were measured in this study; all running-speed assays conducted in this study used horizontal surfaces.

### Data analysis and statistics

To describe the relationship between body mass and running speed, we implemented three complementary analyses. First, we assessed the fit of a recent first-principles biomechanical model that directly predicts the allometry of maximum running speed, using physical parameters estimated from comparative data across the kingdom Animalia [21, 24, 25]. Second, we fitted a version of this model in which muscle mass fraction and the effective coefficient of restitution were allowed to vary, to test whether spiders depart substantially from the trend observed across terrestrial animals. Third, we fitted a conventional allometric power law in log_10_-space. Both fitted models account for phylogeny using PGLS; the predictive biomechanical model uses the independently specified physical parameters without fitting them to the spider data. Quality-of-fit was generally assessed as the root mean-square error (RMSE) of models evaluated on log_10_-transformed speed, to avoid disproportionate weighting of large absolute errors. For the remainder of this manuscript, we refer to these models as (i) predictive biomechanical model; (ii) fitted biomechanical model; and (iii) allometric power law, respectively.

The biomechanical model is based on an energy analysis of an idealised musculoskeletal system and is described in detail elsewhere [21, 24, 25]. For the present study, the key result is that maximum running speed, *v*_max_, is predicted to vary as

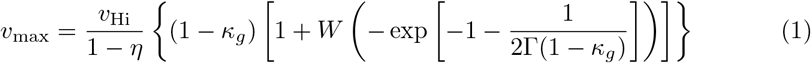

Here, *W* is the principal branch of the Lambert-W function, *v*_Hi_ is the characteristic speed set by maximum muscle shortening velocity and musculoskeletal gearing, *η* is an effective coefficient of restitution, Γ is the physiological similarity index, and *κ_g_*is the reduced parasitic energy associated with gravity. Using a meta-analysis of published data on vertebrate musculoskeletal systems, Labonte et al. [21] estimated 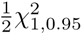, where *m*_kg_ is body mass in kilograms. These values were used for the predictive biomechanical model.

To test whether spiders depart from this estimate, we fitted *η* and the fraction of body mass specialised as leg musculature, *m_f_*, while keeping all other physical quantities at the values used by Labonte et al. [note that *m_f_* is equal to half of the total muscle mass fraction, see 21]: maximum muscle strain rate *ɛ*_max_ = 10s*^−^*^1^, mechanical advantage *G* = 0.3, fascicle-length *l_m_* = (0.03, m) *m*^1*/*3^, muscle density *ρ* = 1060kgm*^−^*^3^, maximum isometric stress *σ*_max_ = 250kPa, and maximum muscle strain *ɛ*_max_ = 0.3. The fitted muscle mass fraction is therefore conditional on those fixed quantities. The governing dimensionless quantities then read,

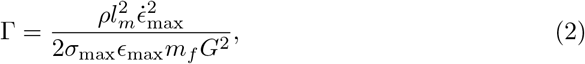

and

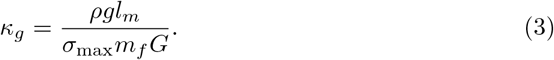

Fitting *m_f_* thus changes both Γ and *κ_g_*, which permits variation in curve shape, whereas fitting *η* controls the velocity scale.

Because species-level observations are not statistically independent, fitted models were estimated using phylogenetic generalised least squares (PGLS; 110–112). In the standard linear case, the expected response is written as *Xβ*, where *X* is the design matrix and *β* is the vector of regression coefficients. For the nonlinear biomechanical model, the residual covariance structure remains unchanged, but the linear predictor *Xβ* is replaced with the model-specific nonlinear mean function *f* (*m, θ*), where *θ* denotes the fitted model parameters, i.e., *η* and *m_f_* [113, 114]. Residuals were therefore defined as *r*(*θ*) = *y − f* (*m, θ*), where *y* = log_10_(*v*), and were evaluated using the phylogenetic covariance matrix *V* . Conditional on *V*, the residual contribution is the phylogenetically weighted residual sum of squares, *r*(*θ*)*^⊤^V ^−^*^1^*r*(*θ*). Thus, the nonlinear analysis differs from standard PGLS only in the form of the mean function and in requiring numerical rather than closed-form parameter estimation.

The phylogenetic covariance matrix was obtained from the pruned tree using the vcv function in the R package ape [115]. We modelled residual phylogenetic signal using Pagel’s *λ* [112, 116], with

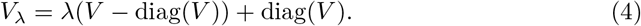

Thus, *λ* = 0 corresponds to phylogenetically independent residuals, whereas *λ* = 1 corresponds to the Brownian-motion covariance implied by the supplied tree.

Model parameters and *λ* were estimated by maximum likelihood. Because the biomechanical model contains bounded parameters and a physical validity boundary at *κ_g_* = 1, we optimised *λ* and *η* on the logit scale, and re-wrote *m_f_* as *m_f,_*_min_ + exp(*θ*), where *m_f,_*_min_ is the smallest value that keeps *κ_g_ <* 1 for the largest observed species. These very features can also make the likelihood surface uneven near the validity boundary and, in principle, allow global shifts in speed magnitude through *η* to be partially confounded with phylogenetically clustered residual deviations, captured by *λ*. To safeguard against these issues, we used multi-start optimisation across *m_f_*, *η*, and *λ*, and assessed local identifiability by computing the Hessian at the maximum-likelihood estimate [113, 114]. The solution was stable across starting values, and the positive-definite Hessian with low condition number indicated a well-resolved local optimum.

For comparison with the fitted biomechanical model, we also fitted the allometric power law,

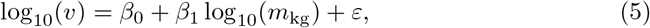

with the same Pagel-*λ* covariance structure. The predictive biomechanical model, fitted biomechanical model, and allometric power law were compared descriptively using RMSE. Because the two fitted models are non-nested, we also report AIC as a secondary comparison [117], but note that we generally do not treat the allometric power law and the fitted biomechanical model as alternative mechanistic hypotheses. In the complete dataset, the main conclusions from subsequent analyses were the same for residuals from the fitted biomechanical model and the allometric power law. To maintain readability, we thus report the results for the fitted biomechanical model in the main text, and the equivalent results for the allometric power law in the supplementary information.

Confidence intervals for fitted parameters were calculated by profile likelihood, following standard practice for nonlinear models and for models with bounded or nuisance parameters such as Pagel’s *λ* [114, 118, 119]. The focal parameter was fixed across a grid of values, the nuisance parameters re-optimised, and the values at which the negative log-likelihood increased by 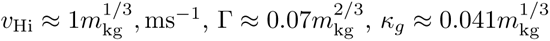 from its minimum identified, following the usual likelihood-ratio approximation [120]. Finally, the mass at which the fitted biomechanical curve reached its maximum was obtained numerically from Eq. 1, and the mass at which predicted speed reaches zero was calculated as the mass for which *κ_g_* = 1. Because *η* rescales the fitted curve but does not alter these critical masses, confidence intervals for both quantities were obtained by propagating the profile-likelihood interval for *m_f_* through the corresponding transformations. Figure 2 shows descriptive uncertainty envelopes spanning the minimum and maximum predictions at each body mass across all combinations of the lower and upper marginal 95% profile-likelihood limits of the fitted mean parameters.

**Fig. 2.**
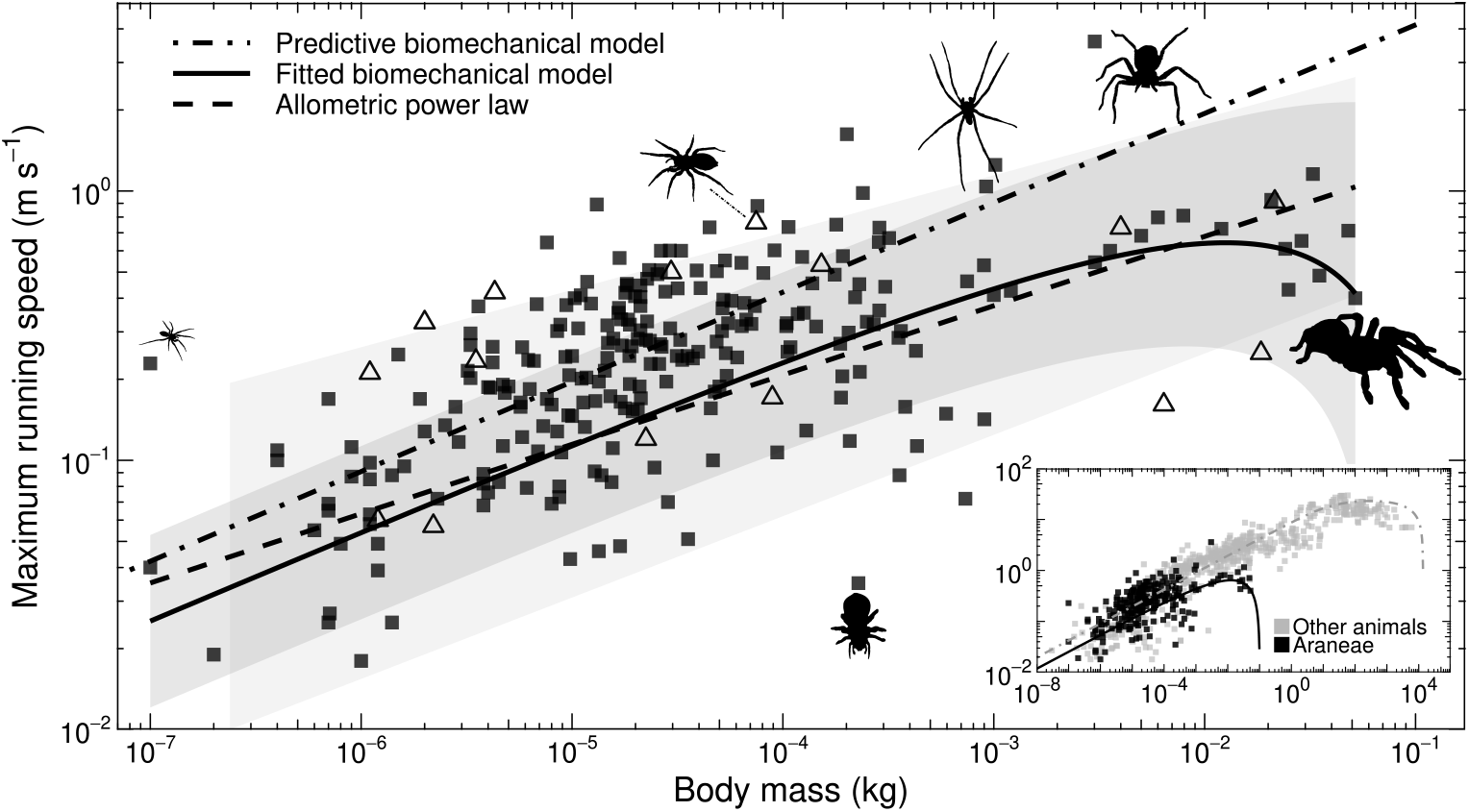
Relationship between maximum running speed and body mass for the fastest recorded individual across 258 spider species. Filled squares indicate species included in the phylogenetic comparative analyses; open triangles indicate species present in the raw dataset but excluded from PGLS analyses because they could not be matched to a terminal taxon in the pruned tree. Across approximately six orders of magnitude in body mass, running speed increased significantly, spanning roughly 0.01–4 m s−1, but comparable variation in speed was also observed among species of similar body mass. Three models were used to describe this relationship: the predictive biomechanical model (dot-dashed line; RMSE = 0.325 log10 m s−1), the fitted biomechanical model (solid line; RMSE = 0.340 log10 m s−1), and the allometric power law (dashed line; RMSE = 0.343 log10 m s−1). Shaded areas show descriptive uncertainty envelopes obtained from marginal 95% profile-likelihood parameter limits for the two fitted PGLS models (see Methods). The inset compares the spider data with a large comparative dataset for other running animals assembled by Labonte et al. 2024 [grey, 21], where the dot-dashed and solid lines are the same as in the main figure.

Body mass explained a substantial share of the variation in maximum running speed, but considerable residual variation remained. To investigate whether this variation was associated with leg morphology, ecological guild, or dominant locomotor orientation, we conducted a series of additional analyses, implemented as extensions of the fitted biomechanical model. Equivalent analyses using the allometric power law are presented in the supplementary information.

To test whether leg morphology explained residual variation in speed, relative leg length and leg slenderness were first log_10_-transformed, mean-centred, and scaled to unit variance. We then fitted

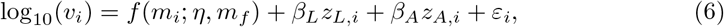

where *z_L_* is relative leg length and *z_A_*is leg slenderness. This model tests whether species with relatively longer or more slender legs run faster or slower than expected from the mass-dependent biomechanical prediction, while accounting for phylogenetic covariance and the other morphology variable. We compared this “morphology model” with the base fitted biomechanical model using a likelihood-ratio test and AIC.

To test whether ecological specialisation or dominant locomotor orientation were associated with speed variation beyond the mass-dependent biomechanical prediction, we fitted separate categorical-offset models. For ecological guild, this model was

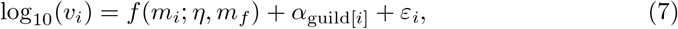

with the same phylogenetic covariance structure as above; an analogous model was fitted for dominant locomotor orientation, coded as upright or inverted. Each model was compared with the base fitted biomechanical model using a likelihood-ratio test, with degrees of freedom equal to the number of additional categorical offsets. Where significant effects were found, pairwise post-hoc contrasts among levels were evaluated using likelihood-ratio tests with Holm correction across all pairwise comparisons. Models including morphology plus guild, and morphology plus locomotor orientation, were also tested as sensitivity analyses; neither categorical predictor significantly improved the morphology model, and these results are therefore not discussed further.

To visualise ecological, locomotor, and morphological effects, we show two types of residual plots. For the categorical predictors, we plotted observed-minus-predicted residuals from the base fitted biomechanical model. These residuals show whether groups tend to lie above or below the mass-dependent biomechanical prediction. For each morphology predictor, *z_j_*, we instead plotted partial residuals, calculated as the full-model residual plus the fitted contribution *β*^^^*_j_ z_j_*. Thus, the partial residual plot for relative leg length, say, shows the association between speed and relative leg length, after accounting for body mass, phylogenetic covariance, and leg slenderness. Positive partial residuals indicate species that run faster than expected after accounting for these terms, whereas negative partial residuals indicate the opposite.

All analyses were conducted in R [121]. Tree handling and phylogenetic covariance matrices used ape [115], ancestral-state and phylogenetic visualisations used phytools where required [122], and standard linear/mixed-model routines used nlme where applicable [123]. The nonlinear phylogenetic likelihood for the biomechanical model was custom-written, because standard PGLS routines assume a linear mean function.

## Results

### Allometry of running speed

Across 258 species from 68 out of 139 extant spider families, running speed increased substantially with body mass, from a minimum of 0.018 m/s measured for the money spider *Maso sundevalli* (body mass 1 mg), to a maximum of 3.59 m s*^−^*^1^ recorded for the huntsman spider *Heteropoda cervina/jugulans* (body mass 3.01 g; see Fig. 2). Notably, running speeds for several of the 28 species with a body mass above 0.5 g seemed to fall below a simple allometric trendline (see below), dropping as far as 0.4 m/s for the heaviest spider included in this dataset (Salmon Pink Birdeater -*Lasiodora parahybana*, 51.8 g).

The predictive biomechanical model yielded an RMSE of 0.325 log_10_(m s*^−^*^1^), evaluated for the 243 of the 258 species for which phylogenetic data were available, to make results comparable. The fitted biomechanical model had a similar RMSE of 0.340 log_10_(m s*^−^*^1^) and AIC = 35.46, with fitted parameter estimates of *m_f_* = 0.00195 (95% CI [0.00167, 0.00316]) and *η* = 0.82 (95% CI [0.62, 0.91]); residuals showed strong phylogenetic signal (*λ* = 0.82, 95% CI [0.68, 0.90]). The alternative, based on a classic allometric power law, yielded log_10_(*v*_max_) = 0.35 + 0.26 log_10_(*m*), where *m* is body mass in kilograms; 95% confidence intervals were [0.001, 0.697] for the intercept and [0.21, 0.30] for the slope. This model had a nearly identical RMSE of 0.343 log_10_(m s*^−^*^1^) and slightly higher AIC = 37.39; its residuals likewise retained a similar phylogenetic signal (*λ* = 0.79, 95% CI [0.62, 0.89]).

Although body mass explained a significant fraction of variation in running speed, substantial variation remained even among spiders of very similar body mass. For example, the best performing individuals of the orange purse-web spider, *Calommata signata*, and the tube-web spider, *Segestria florentina*, both weighed approximately 200 mg, yet their maximum running speeds differed by 28-fold—not far off the speed variation of the dataset as a whole (Fig. 2).

### Morphological and ecological correlates of running speed

Leg and carapace dimensions were strongly correlated with body mass, scaled close to isometry (Fig. 3), and showed significant phylogenetic signal. Leg I length scaled with body mass with a PGLS slope of 0.38 (95% CI [0.35, 0.42]; PGLS *R*^2^ = 0.71; *λ* = 0.92, 95% CI [0.82, 0.97]), and leg IV length with a slope of 0.37 (95% CI [0.34, 0.40]; PGLS *R*^2^ = 0.79; *λ* = 0.93, 95% CI [0.82, 0.98]). Carapace area scaled with a slope of 0.64 (95% CI [0.61, 0.68]; PGLS *R*^2^ = 0.96; *λ* = 0.48, 95% CI [0.19, 0.76]). Thus, there was no evidence for substantial systematic changes in body shape with body size, and we instead focused on the variation in size-corrected speed explained by size-specific shape variation.

**Fig. 3.**
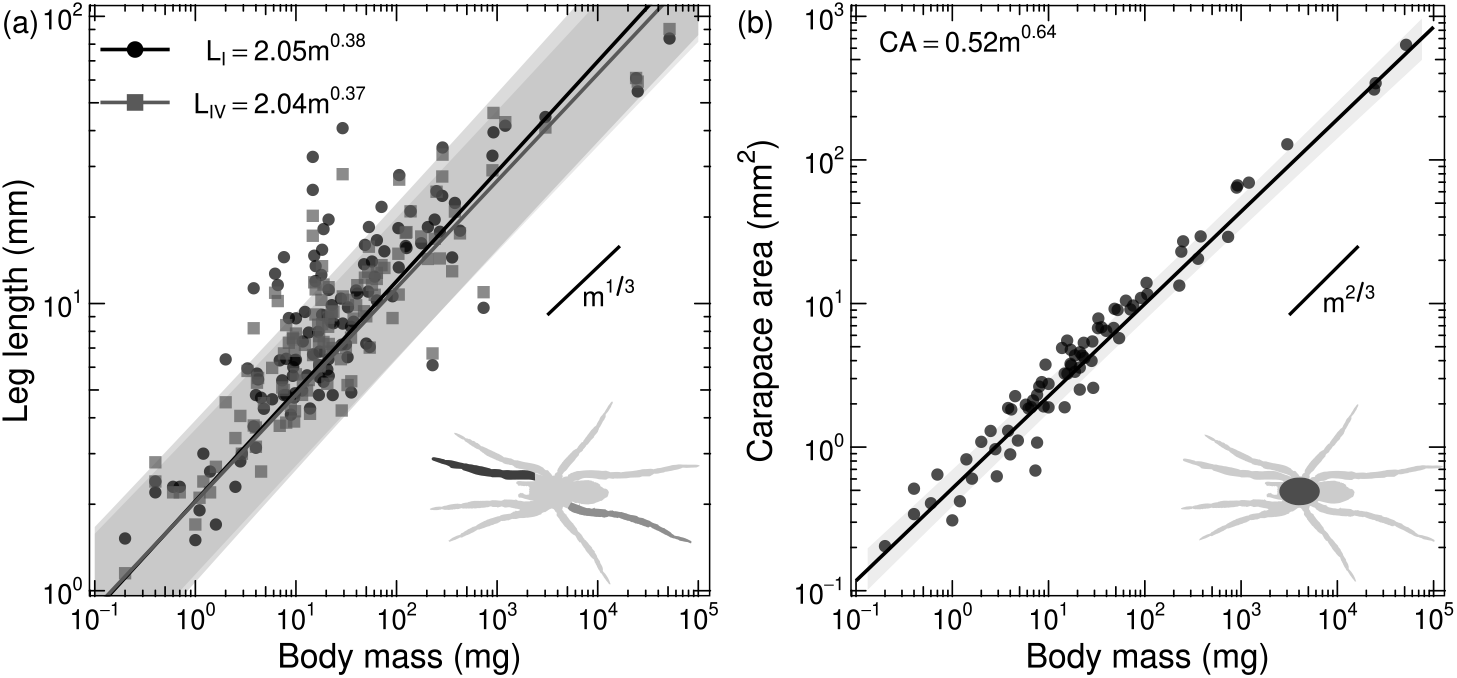
Scaling of leg lengths and carapace area with body mass. Panel (a) shows leg I length (LI) and leg IV length (LIV) for 123 species; panel (b) shows carapace area for 78 species, calculated as the area of an ellipse from carapace length and width. Points represent species-level measurements, and fitted lines show phylogenetic generalised least-squares regressions on log10-transformed data. Shaded bands indicate pointwise 95% profile-likelihood confidence intervals for the fitted regressions. Short line segments show the parsimonious isometric expectation: slope 1/3 for linear dimensions and 2/3 for area. All three traits scaled close to isometry and showed significant phylogenetic signal.

Adding relative leg length and leg slenderness to the fitted biomechanical model significantly improved model fit on the morphology-complete subset (*n* = 157; likelihood-ratio test: LR = 29.56, df = 2, p *<* 0.001): RMSE decreased from 0.344 to 0.292 log_10_(m s*^−^*^1^), and AIC decreased from 39.44 to 13.88. This improvement was driven primarily by relative leg length (Fig. 4 a), which had an estimated effect of 0.099 log_10_(m s*^−^*^1^) per standard deviation increase in log_10_ relative leg length (95% CI [0.012, 0.188]), corresponding to an approximately 26% increase in running speed (95% CI [2.7%, 54%]). The effect of leg slenderness was also positive, but weaker and statistically uncertain (Fig. 4 b. Estimated effect = 0.041 log_10_(m s*^−^*^1^) per SD, 95% CI [-0.047, 0.129]). Adding guild to the morphology model did not improve fit (LR = 2.30, df = 5, p = 0.81; AIC increased from 13.88 to 21.57), and neither did adding dominant locomotor orientation (LR = 0.002, df = 1, p = 0.97; AIC increased from 13.88 to 15.88).

**Fig. 4.**
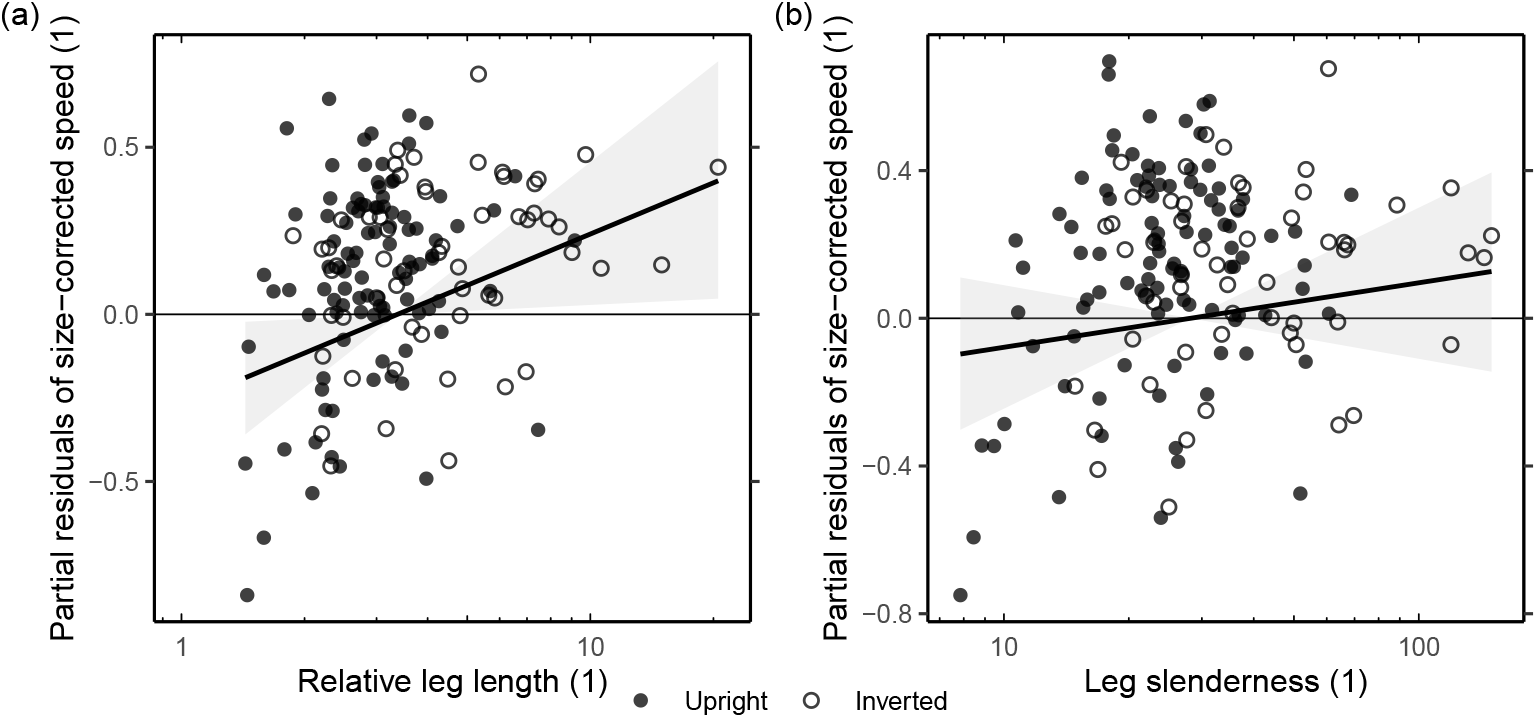
Partial residual plots showing the association between size-corrected running speed and leg proportions. Panel (a) shows relative leg length, calculated as mean leg I and IV length divided by carapace length; panel (b) shows leg slenderness, calculated as mean leg I and IV length divided by mean tibia I and IV width. Closed symbols indicate species with predominantly upright locomotion, and open symbols indicate species with predominantly inverted locomotion. Morphological predictors were fitted as log10-transformed, mean-centred, and scaled variables, but are plotted here on their original ratio scale. Partial residuals were calculated from the full morphology model: the relative-leg-length effect is shown after accounting for body mass, phylogenetic covariance, and leg slenderness, whereas the leg-slenderness effect is shown after accounting for body mass, phylogenetic covariance, and relative leg length. Lines show the fitted partial effects from the fitted biomechanical model with morphology, with shaded areas indicating pointwise 95% profile-likelihood uncertainty envelopes. Relative leg length had a significant positive effect on size-corrected running speed, whereas the effect of leg slenderness was positive but statistically uncertain.

Across the full phylogenetically matched dataset (*n* = 243), ecological guild explained a smaller, but statistically supported, fraction of residual variation in running speed (Figs. 5 & 6). Adding guild-specific offsets to the fitted biomechanical model improved fit relative to the base model (likelihood-ratio test: LR = 12.76, df = 5, p = 0.026), reduced RMSE from 0.340 to 0.296 log_10_(m s*^−^*^1^), and decreased AIC from 35.46 to 32.71. The largest positive fitted offset occurred in ground-active hunters (guild 4; offset = 0.287 log_10_(m s*^−^*^1^), 95% CI [0.016, 0.559]), corresponding to about two-fold increase in speed over the reference guild (95% CI [1.04, 3.62]). Ground-active hunters also had the highest mean residual speed, running on average about two-fold faster than predicted by the base fitted biomechanical model (mean residual = 0.306 log_10_(m s*^−^*^1^), 95% CI [0.222, 0.389]). By contrast, burrow/tube-web builders (mean residual = -0.011, 95% CI [-0.193, 0.172]) and ambush hunters (mean residual = -0.056, 95% CI [-0.263, 0.151]) were centred close to, or slightly below, the fitted expectation (Fig. 5). No pairwise guild contrast remained significant after Holm correction (see supplementary information), so these group-level differences should be interpreted as the main contributors to the global guild effect rather than as individually resolved pair-wise differences. Last, dominant locomotor orientation did not improve the base fitted biomechanical model (LR = 0.85, df = 1, p = 0.36; AIC increased from 35.46 to 36.62).

**Fig. 5.**
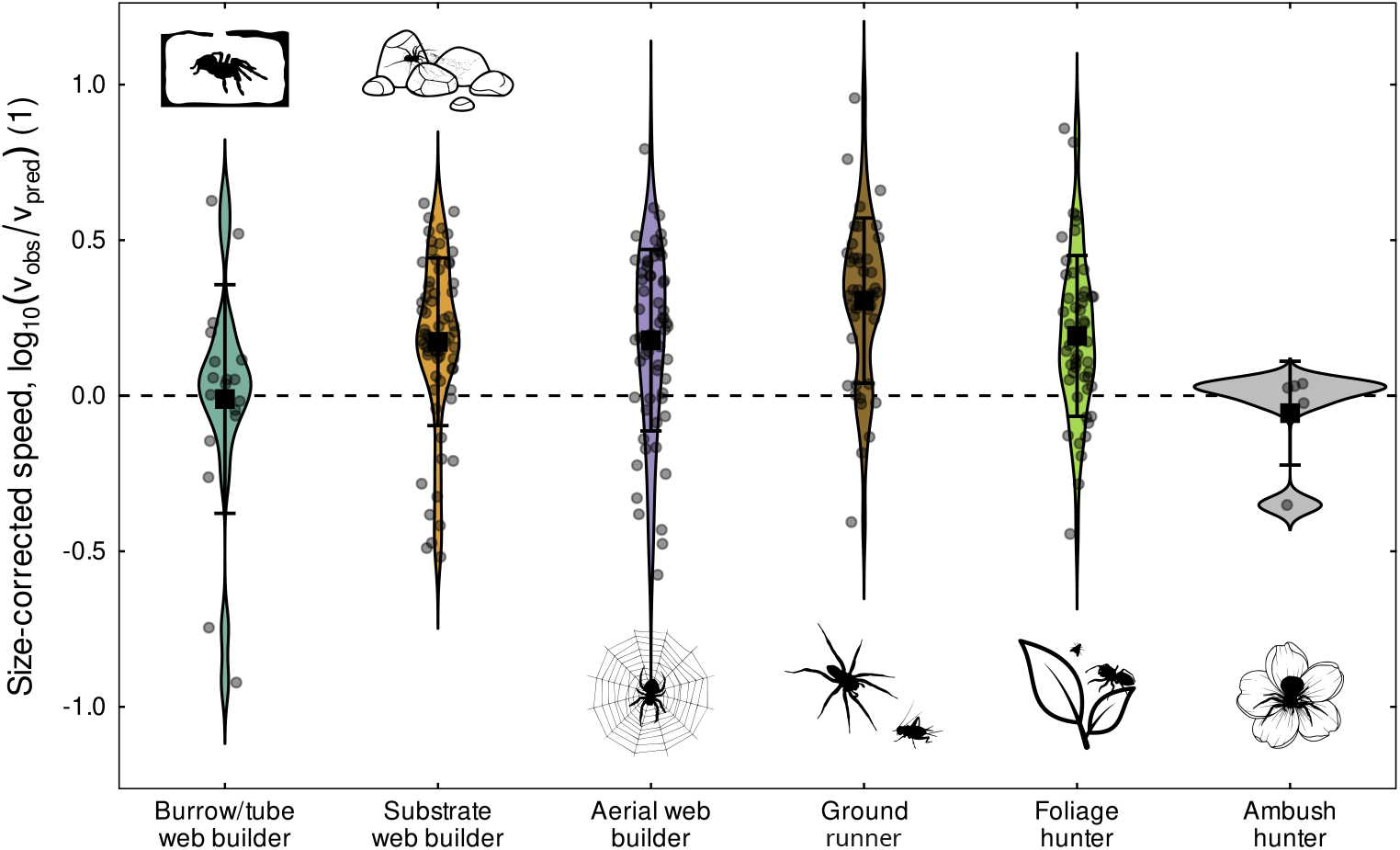
Distribution of size-corrected maximum running speed across ecological guilds, shown as logarithmic residuals from the fitted biomechanical model (solid line in Fig. 2). Violin plots show residual distributions; grey points are species, black squares are guild means, and error bars show standard deviations. Guild explained a significant component of residual variation overall, although no pairwise guild contrast remained significant after Holm correction. Ground-active hunters tended to run faster than expected for their size, whereas ambush hunters showed the slowest size-corrected speeds on average, and the narrowest residual spread.

**Fig. 6.**
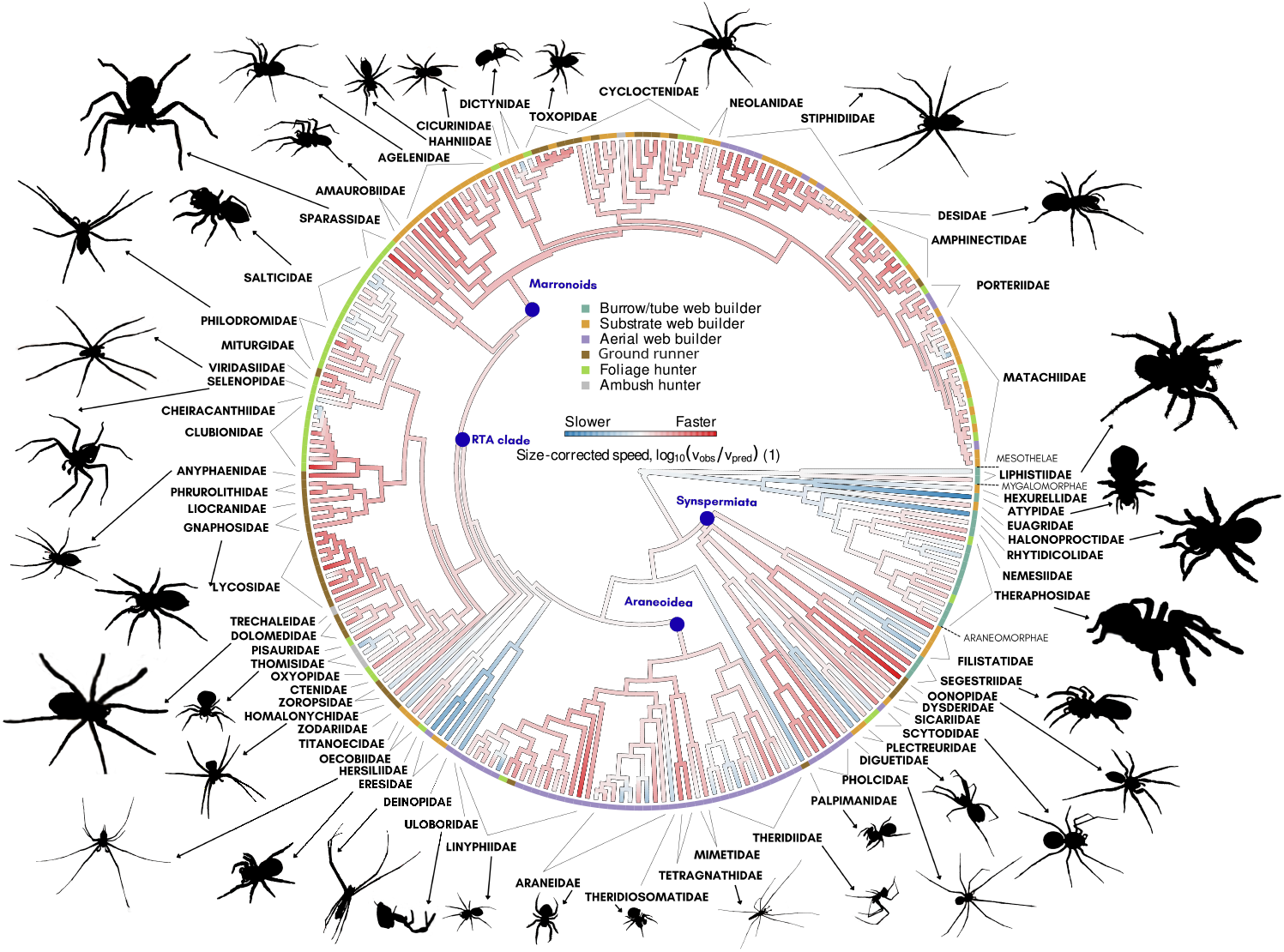
Brownian-motion based ancestral character estimation of size-corrected running speed across the spider phylogeny, estimated from the logarithmic residuals from the fitted biomechanical model (solid line in Fig. 2). The reconstruction suggests that early-branching, heavy-bodied lineages remained close to the running speeds expected for their size, whereas araneomorphs diversified markedly, with some lineages running more quickly and some more slowly than expected from their size alone. The blue dots mark the most recent common ancestors of the sampled RTA clade, Marronoids, Araneoidea and Synspermiata. In particular, the evolutionarily younger RTA-clade tends towards higher relative running performance.

### Macroevolutionary patterns

Visualisation of the phylogenetic structure in size-corrected running speed via ancestral state estimation suggested that representatives of the early-branching, heavy-bodied lineages—mygalomorphs and liphistiids—run about as fast as expected for their size. Variation among araneomorphs, in turn, is much greater, with some lineages tending to run much faster or much slower than expected from their size alone (Fig. 6). For example, within the Synspermiata clade, Dysderoidea and Pholcidae exhibited higher size-corrected running speeds than their corresponding sister lineages, independent of ecological differentiations within these groups. Among Araneoidea, a large clade of predominantly aerial web builders, size-corrected running speed also varied substantially. Palpimanidae, Eresidae, Deinopidae, and Uloboridae consistently exhibited evolutionary trends towards running speeds lower than predicted from mass alone, whereas most representatives of the evolutionarily younger RTA-clade run faster than expected for their size. Within the RTA-clade, a tendency towards reduced size-corrected running speeds occurred in Thomisidae (crab spiders) and Salticidae (jumping spiders).

## Discussion

By combining comparative phylogenetic methods with biomechanical modelling, this study examines how variation in spider running speed emerges from the interplay of body size, morphology, phylogenetic history, and ecology. Across a broad and taxonomically diverse sample, we found substantial size-related variation in maximum running speed, but also that much of the biologically meaningful variation emerges only once performance is considered relative to size: size-corrected running speed was associated with relative leg length and guild, showed clear phylogenetic structure, but was not consistently associated with preferred locomotor orientation or leg slenderness. Below, we discuss each of these factors in turn and consider how the present results inform broader ideas about the evolution of running performance in arthropods.

### The allometry of maximum running speed in Araneae: complex or simple?

Running speed is an unusual performance trait because its relationship with body size is non-monotonic: the fastest animals are neither the largest nor the smallest, but of intermediate size [19]. Recently, this unusual allometric pattern was linked to size-dependent variation in two distinct skeletal-muscle energy capacities, and to an increasing importance of gravitational potential energy in larger animals [21, 24]. A direct prediction from this biomechanical model described the data about as well as a conventional allometric power law (Fig. 2). Spiders, to first order, thus run about as fast as expected for their size, and body size—across many orders of magnitude—is a robust predictor of running performance.

This is not to say, however, that all meaningful information is contained in but one scaling law. The fitted biomechanical model revealed a curious deviation. The original estimate of the effective coefficient of restitution, *η* = 0.89, remained within the 95% confidence interval of the spider-specific fit (*η* = 0.818, 95% CI [0.621, 0.913]). The fitted muscle mass fraction, by contrast, was about a factor of 50 lower than the value used in the predictive biomechanical model (*m_f_* = 0.19%, 95% CI [0.17, 0.32%], compared with 10%). As a result, the spider-specific fit predicts a peak in running performance at an intermediate body mass of 12.6 g (95% CI [8.1, 50.3] g), and a zero-speed limit at 103 g (95% CI [65, 437] g). Spiders, in other words, are predicted to replicate at a lower mass range the complex allometric pattern observed across the kingdom Animalia at large.

Our data appear visually consistent with a reduction in absolute running performance in the heaviest spiders (Fig. 2), which has previously been suggested to result from the increasing difficulty of lifting bulky bodies against gravity [26, 34]—exactly the physical effect that accounts for the drop in speed in the biomechanical model [21]. Nevertheless, the statistical evidence for such a drop is weak: the allometric power law fits the data about as well but predicts a monotonic change. The poor statistical power is, in part, an issue of data distribution. There are far fewer heavy than lightweight spiders, and least-squares fits are therefore dominated by the pattern in smaller spiders, where both models predict a similar increase in speed. A statistical distinction is there-fore challenging and will only be possible through a concerted effort to increase the sample size for heavy spiders. The biomechanical model offers a complementary test: independently measured or estimated physiological parameters can establish whether a downturn is physically plausible.

Qualitatively, a reduction in the body-mass fraction allocated to limb musculature may seem surprising. Why should it occur? Here, the biomechanical model is instructive: in small animals, maximum running performance is predicted to be governed to first order by muscle length, maximum strain rate, and gearing; in other words, it is independent of muscle mass [21, 24]. On this basis, previous work predicted that selection for large muscle mass may be relaxed at small body sizes, producing a repeated “hump” in the speed–mass relationship [21]. Preliminary support for this hypothesis exists: reptiles have a muscle mass fraction roughly half that of mammals and appear to slow down at a lower body mass, both across and within species [21, 124, 125]. The present results suggest that this pattern may be replicated once again in spiders, at a smaller scale—a phenomenon reminiscent of what Meunier termed “transpositional allometry” [126–128], and which appears repeatedly in biomechanical scaling data [129–131].

Quantitatively, however, great scepticism is warranted: a muscle mass fraction of about 0.2% is so small as to seem arguably implausible altogether. But the muscle mass fraction was selected as just one possible parameter capable of shifting the curve shape, precisely because there was an *a priori* reason to expect it might vary; other physical quantities may differ between spiders and other terrestrial runners, too. Concretely, the speed prediction is preserved for any combination of parameters that leaves unchanged the velocity scale *v*_Hi_*/*(1 *− η*), the physiological similarity index Γ, and the reduced parasitic force *κ_g_*. To give but one example of what this could look like in practice, if spiders shortened their muscles by about 20 rather than 30%, had a representative mechanical advantage of *G ≈* 0.02 rather than 0.3, a maximum muscle strain rate of *ɛ*_max_ *≈* 2.1 s*^−^*^1^ rather than 10 s*^−^*^1^, and an effective coefficient of restitution of *η ≈* 0.42 rather than 0.89, a much more plausible muscle mass fraction of 3% would yield the same fitted curve. These values are all physically plausible, and a lower mechanical advantage is, in fact, even expected in smaller animals, both empirically and theoretically [132, 133]. Direct experimental measurements of these quantities across a robust comparative sample of spiders are needed to bring certainty and provide a robust test for the predictive accuracy of the biomechanical model.

Of course, another limitation of the biomechanical model has been hiding in plain sight: spiders do not rely solely on direct muscle actuation. Instead, hydraulic pressure is used to extend several major joints [29, 30]. The implications of this mixed actuation are addressed in the next section.

### Hydraulics and the role of leg proportions

The biomechanical model is built on the assumption that all movements are driven directly by muscle contraction. This assumption is appropriate for many animals, but it is incorrect for spiders, in which leg extension can be driven by hydraulic actuation via haemolymph flow instead [29, 134, 135]. This pressure, elevated during activity by contractions of body musculature [134, 136], stiffens the leg arch, which then acts as an antagonist to the flexor muscles, especially at the femur–patella and tibia–metatarsus joints, which lack extensor muscles [30, 137–140]. Might speed, then, be limited by the performance of the hydraulic system [34, 141, 142]? Two lines of evidence speak against this conclusion.

First, previous experimental work on the large wandering spider *Ancylometes concolor* concluded that body propulsion is powered primarily by leg flexion, combined with vertical impulses supplied by rotation at the trochanter–femur joint—movements that are all muscle-driven [30]. In the same vein, it has been hypothesised that, if the hydraulic leg extension was limiting peak running speed in large spiders, leg extension should be slower than leg flexion [34]. However, allometric data on two araneomorph spiders showed that, if anything, the opposite is the case [34]. Hydraulic actuation could still impose costs indirectly, for example by increasing the work required for leg flexion. But here, too, recent theoretical work on haemolymph flow through the leg lacunae suggested that such a limitation is unlikely to significantly reduce muscle performance [143].

Second, if whole-leg dimensions reflected the relevant hydraulic flow paths, longer and narrower legs might be expected to incur greater flow resistance and limit speed. However, our data, which include several long-legged taxa such as Pholcidae and Tetragnathidae, suggest the exact opposite (Fig. 4 a): both relative leg length and leg slenderness were associated with an increase in size-specific running performance, significantly so for relative leg length. The sum of the available evidence thus suggests that the hydraulic system—which ultimately relies on muscle contraction, too—is at least not obviously imposing a limit independent of that imposed by muscle physiology. Detailed work, in particular in small spiders, where our understanding of hydraulics is more limited, and including more comprehensive and accurate geometric models of flow pathways and estimates of haemolymph viscosity, is needed to place these preliminary conclusions on a robust footing.

The positive effect of relative leg length is interesting also from an ecomechanical perspective, for it appears a core locus of ecological specialisation for high running speeds: guild no longer had a significant effect once variation in leg proportions was accounted for. Relatively longer limbs were consistently related to elevated running performance also in lizards [144–149], but this pattern is still in want of a comprehensive mechanistic explanation. The classic argument is that longer legs permit longer strides, and thus, at equal stride frequency, faster speeds. This is surely correct, but it also tacitly assumes that muscle can accommodate the increased work and power demand that comes with increasing stride lengths at constant stride frequency. One may speculate that this demand is, in some sense, met automatically, because longer legs are likely to house longer flexor muscles, which allow faster shortening speeds, and thus bring with them a higher kinetic energy capacity—the likely limiting element of movement at small scales [20, 21, 25, 133]. But this is not necessarily so, and an increase in limb length may equally well be associated with a disproportionate growth of apodemes instead.

Even though the magnitude of the effect of relative leg length was not trivial—across the full observed range of relative leg lengths, it accounts for an approximately four-fold difference in speed—it does not account for all variation that remains after body mass is accounted for. This is, no doubt, in part due to measurement error, because maximum performance is difficult to elicit. But it may also reflect other morphological differences that our reduced set of morphological characters did not capture, including the size and architecture of leg and prosomal musculature, relative size differences between hind and front legs [28], or the angular orientation of leg flexion planes [150]. Laterigrade legs, i.e. morphologies where the leg flexion plane is tilted from near vertical to near horizontal, have evolved multiple times in araneomorph spiders, presumably because they enhance manoeuvrability [150]. Some species with laterigrade or near-laterigrade legs in our dataset, such as *Heteropoda* spp., *Toxops montanus*, *Toxopsoides* spp. and *Manjala plana*, indeed exhibited elevated size-specific speeds, but others, such as *Philodromus spp.*, *Selenops*sp. and *Eusparassus dufouri* did not. Thus, even within families (Sparassidae), the tradeoff between manoeuvrability and running speed may not be straightforward.

### Does running performance evolve as an adaptation to hunting mode?

Running is an important behaviour in spiders, underpinning hunting, dispersal, mate search, and the evasion of predators [151, 152]. It is therefore reasonable to expect running performance to have evolved in concert with behavioural niches and microhabitat preferences.

Consistent with this expectation, the inclusion of guild improved the biomechanical model fit. Ground-dwelling cursorial (active pursuit) spiders had the highest mean size-corrected speeds, whereas tube-web and burrowing spiders and web-less ambushing spiders tended to exhibit lower size-corrected speeds—arguably as one may expect from the corresponding foraging strategies (Fig. 5). In addition, there were repeated evolutionary changes in size-corrected running speed that coincided with shifts in ecological niche (Fig. 6). Within the large infraorder Araneomorphae, for example, the aerial web-building Pholcidae were faster than their sister lineage within Synspermiata, the substrate-sheet-weaving Plectreuridae—a difference that is also reflected in divergent trends in leg length [28]. These effects, however, were modest overall; pairwise guild comparisons of the mass-corrected running speed did not show significant differences, and there were several instances where ecological divergence was not accompanied by a shift in size-corrected running speed (Fig. 6). Within Dysderoidea, for example, the ground-running Dysderidae and Oonopidae run much faster than expected for their size -but so do the tube-web-dwelling Segestriidae. Given that tube webs probably evolved earlier and cursorial lifestyles are derived [153], this similarity is unlikely to be explained by evolutionary lag but instead suggests that elevated running speeds evolved before the shift to a free-hunting lifestyle in Dysderoidea. Such changes in locomotor performance may partially be explained with changes in the respiratory system. The older lineages of Mesothelae, Mygalomorphae and some early diverging Araneomorphae exhibit two pairs of book lungs, but no tracheae [154]. In contrast, most araneomorphs have only one pair of book lungs and, in addition, tracheae. Dysderoidea and many representatives of the RTA-clade exhibit extensive tracheae, which extend into the prosoma and legs and may allow better oxygenation of muscles [154, 155].

One possible explanation for the generally weak association between size-corrected running performance and guild is that the broad guild categories used in this study capture only part of the fine-grained variation in hunting style relevant to running performance. In support of this view, both Thomisidae (crab spiders) and Salticidae (jumping spiders) tended to run more slowly than expected for their size, likely reflecting specialisation for ambush predation (in the Thomisidae) and prey capture through targeted jumps (in Salticidae). Notably, this difference is maintained even in the highly derived morphology exhibited by the ant-mimic *Synageles venator*. More broadly, the shape of the adaptive landscape of running speed is surely affected by ecological factors outside specific prey capture strategies, such as microhabitat structure. For example, in arboreal habitats, species that live in fine-structured foliage may rely on manoeuvrability more than raw speed. And last, the relation between ecological guild and locomotor performance may be weakened by sexual dimorphism and the diverging ecological requirements of males and females [41, 47, 156]. Even where extreme dimorphism is absent, males are often more mobile than females and may differ systematically in body proportions relevant to locomotion. This can be observed in active hunters such as Lycosidae, where the males are smaller, have relatively longer legs, and a wider cephalothorax [39]; and in some subterranean and sedentary web-building taxa, males can have more cursorial body shapes, presumably to help with mate search [40, 46, 47]. Another noteworthy result is that the contrast between the largely non-web-building RTA clade and the predominantly aerial-web-building Araneoidea was not reflected in strongly divergent evolutionary patterns (Fig. 6), in seeming contradiction with both the gravity hypothesis [26], and the trait substitution hypothesis [31].

The gravity hypothesis purports that adaptation to suspensory locomotion in aerial webs favours longer, thinner legs, and that these would reduce locomotor performance on horizontal surfaces, because they are less well suited for weight support [26]. This idea can be read in two ways: first, as a prediction primarily about morphology, and second, as a prediction about running performance more directly. The morphological prediction is broadly supported: both orb- and cob-web spiders, for example, tend to have longer and more slender legs than ground-dwelling cursorial species [26, 28]. The performance prediction, however, is not. In our dataset, higher relative leg length was positively associated with size-corrected running speed (Fig. 4 a). Likewise, species moving predominantly in an inverted, suspensory position in their webs did not differ in size-corrected running speed from species moving mainly upright on the substrate. Perhaps, as noted by Weihmann [43], simple inverted-pendulum models, on which the gravity hypothesis partly relies, do not lend themselves to application in running spiders, which exhibit only limited vertical oscillations of the body’s centre of mass during locomotion.

The trait-substitution hypothesis offers a different suggestion, namely that the use of snares reduces selective pressure on locomotor performance because prey capture is partly outsourced to the web [31]. Our results provide little support for this idea either. Pairwise comparisons did not show significant differences in size-corrected running speed between cursorial and web-building guilds (Fig. 5), and Kelly et al. [31] reported a similar pattern in marronoid spiders, a clade that shows repeated transitions between web-building and cursorial lifestyles, and which is also represented in our dataset.

## Conclusion

Performance is a key contributor to fitness and is therefore a target of selection, which operates within the bounds of physical laws, and is curtailed further by phylogenetic constraint. Untangling the relative contributions of these factors to phenotypic diversity is as important for our understanding of macroevolutionary processes as it is challenging in practice. Here, we attempted such a deconstruction by combining a large comparative performance dataset with phylogenetic comparative methods and biomechanical analyses. Our results suggest that ecology, phylogenetic history, and size-related variations in biomechanical capacity all contribute meaningfully to differences in peak running performance across the Araneae, and that an integrated approach can help render such qualitative statements increasingly quantitative. In an age in which machine learning and artificial intelligence are coming to dominate research across the sciences, continued progress will depend above all on large-scale comparative experimental work—spanning anatomy, physiology, ecology, and behavioural assays. Only with such data at hand will we be able to eventually drive major advances in our understanding of the factors that shape locomotor performance in spiders, other arthropods, and animals more generally.

## Declarations

Ethics approval and consent to participate Not applicable.

## Consent for publication

Confirmed.

Availability of data and materials The data supporting this study are provided in the Supplementary Information.

## Authors’ contributions

SK collected most of the specimens, running speed and morphometry data, and led data analysis. EB helped with initial morphometry measurements, and VJ contributed to the recording of spiders. HB helped with camera tracking, recording and analysing data from tarantulas on the treadmill. JOW contributed to supervision, specimen collection, and data analysis. MK helped with spider collection, performed the molecular lab work, assembled the molecular matrix and performed the phylogenetic inference. DL conceived of and supervised the project, provided theoretical background and code for biomechanical analysis, and key laboratory resources. The manuscript was written by SK, JOW, and DL. All authors read and approved the final version.

## Supporting information

Raw data

## Acknowledgements

We gratefully acknowledge members of the Evolutionary Biomechanics Laboratory at Imperial College London for advice, technical support, and spider husbandry, with special thanks to Fabian Plum for help with OmniTrax. We thank Siddharth Kulkarni and Gustavo Hormiga for providing most of the UCE contigs that helped stabilise the phylogenetic backbone; Maitry Jani, Susan Kennedy, and Henrik Krehenwinkel for amplicon labwork and sequencing, de novo assembly from fresh specimens; and Daniele Liprandi and Tom Illing for data management, curation, and troubleshooting analyses on the UG brain cluster. Further, we thank Milena Zerbe and Ronja Eilhardt for their help with photographing and taking morphometric measurements of spiders. We are grateful to the management of the London Wildlife Trust and Perivale Wood, Liz Andrew and Adrian Brooker of Hampstead Heath, Cameron Findlay of Bedfont Lakes, Alice Laughton and Rachel Harris of Holland Park, and Darren Hill of Frensham Common, for granting permission to collect spiders; and to Samuel Fabian, Jason Steel, and Lloyd Davies for providing specimens. For tarantula recordings, we thank Gen Popovici, Ray Hale of the British Tarantula Society, and Mark and Luke from SpaSpiders for arranging access to numerous species from their collections. We also thank Frederik Leck Fischer, William Mason, Arno Grabolle and Jim McLean for permitting us to use their photographs to prepare drawings in diagrams. We further acknowledge Esmond Brown for helping with the collection of numerous specimens, identifying multiple species, and contributing images for figures, and Simeon Indzhov for consistently valuable guidance on spider identification and biology. We also thank Dr. Irem Sepil for help with anaesthetising spiders for morphometric measurements. JOW thanks Braxton Jones, Marshal Hedin, Raymond Wyatt Mendez, Monica Sheffer, Jacob Gorneau, Lauren Esposito, Pedro Cardoso, Sergio da Silva Henriques, Arie van der Meijden, and Milan R^̌^eźc for help with spider collection and fieldwork logistics, and Paula Heinz, Josefine Kreuz, and John Seifert for help with animal maintenance.

## Supplementary information: Evolutionary biomechanics of maximum running speed in spiders (Araneae)

### Supplementary information

Repeating the residual analyses using the log-linear allometric PGLS model gave results closely matching those obtained with the biomechanical model.

Adding relative leg length and leg slenderness significantly improved model fit relative to the base allometric PGLS model (likelihood-ratio test: LR = 34.48, df = 2, p *<* 0.001). RMSE decreased from 0.358 to 0.294 log_10_(m s*^−^*^1^), and AIC decreased from 41.61 to 11.13. As in the biomechanical analysis, this improvement was driven primarily by relative leg length, which had an estimated effect of 0.101 log_10_(m s*^−^*^1^) per standard deviation increase in log_10_ relative leg length (95% CI [0.013, 0.190]). This corresponds to an approximately 26% increase in predicted running speed per standard deviation increase in relative leg length (95% CI [3%, 55%]). The effect of leg slenderness was again weaker and uncertain (estimated effect = 0.051 log_10_(m s*^−^*^1^) per SD, 95% CI [-0.037, 0.137]), corresponding to an estimated 12% increase in speed, but with confidence intervals spanning both negative and positive effects (95% CI [-8%, 37%]).

Categorical predictors also gave results similar to those from the biomechanical model. Adding ecological guild to the allometric PGLS model significantly improved fit relative to the base allometric model (likelihood-ratio test: LR = 13.25, df = 5, p = 0.021), reducing RMSE from 0.343 to 0.300 log_10_(m s*^−^*^1^) and decreasing AIC from 37.39 to 34.13. Ground-active hunters again had one of the largest positive fitted offsets (guild 4; offset = 0.271 log_10_(m s*^−^*^1^), 95% CI [0.005, 0.541]), corresponding to an estimated 1.87-fold higher speed than the reference guild (95% CI [1.01, 3.48]). Ambush hunters had the lowest fitted offset (offset = -0.081, 95% CI [-0.435, 0.277]). However, as before, no pairwise guild contrast remained significant after Holm correction, so these differences should be interpreted as contributors to the global guild effect rather than individually resolved pairwise differences.

Dominant locomotor orientation did not improve the base allometric model (LR = 0.86, p = 0.35; AIC increased from 37.39 to 38.53). Adding locomotor orientation after morphology also did not improve fit (LR = 0.16, p = 0.69; AIC increased from 11.13 to 12.97), and had little effect on the morphology coefficients.

In a nutshell, all major conclusions reported in the main text were insensitive to whether body size was accounted for using the biomechanical model or a conventional allometric PGLS model.

Exclusion of adult males left 207 species (132 with complete morphology) and retained the qualitative conclusions. In the biomechanical and allometric models, respectively, relative-leg-length effects were 0.089 (95% CI [0.005, 0.174]) and 0.090 (95% CI [0.006, 0.175]) log_10_(m s*^−^*^1^) per SD of log_10_ relative leg length, and guild effects remained supported (p = 0.036 and 0.043). Leg-slenderness confidence intervals included zero, and locomotor orientation remained unsupported (p = 0.335 and 0.366). Neither guild nor orientation improved fit after morphology (all p *>* 0.79).

**Table 1.** Pairwise comparisons among ecological guilds for residual running speed under the nonlinear biomechanical PGLS model. Contrasts are from likelihood-ratio tests comparing guild-offset models, with Holm correction applied across all pairwise guild comparisons. The log_10_ contrast is defined as guild A minus guild B; speed ratios are therefore 10^Δ^. No pairwise contrast remained significant after Holm correction.

| Guild comparison | Δ log<sub>10</sub>(*v*) | Speed ratio | LR | *p* | *p*<sub>Holm</sub> |
|---|---|---|---|---|---|
| 1–2 | -0.182 | 0.66 | 1.86 | 0.173 | 1.000 |
| 1–3 | -0.282 | 0.52 | 3.93 | 0.047 | 0.520 |
| 1–4 | -0.287 | 0.52 | 4.32 | 0.038 | 0.491 |
| 1–5 | -0.221 | 0.60 | 2.79 | 0.095 | 0.850 |
| 1–6 | 0.050 | 1.12 | 0.08 | 0.783 | 1.000 |
| 2–3 | -0.100 | 0.79 | 2.54 | 0.111 | 0.850 |
| 2–4 | -0.105 | 0.78 | 2.85 | 0.091 | 0.850 |
| 2–5 | -0.038 | 0.92 | 0.47 | 0.492 | 1.000 |
| 2–6 | 0.232 | 1.71 | 2.97 | 0.085 | 0.850 |
| 3–4 | -0.005 | 0.99 | 0.00 | 0.945 | 1.000 |
| 3–5 | 0.062 | 1.15 | 0.72 | 0.395 | 1.000 |
| 3–6 | 0.332 | 2.15 | 5.34 | 0.021 | 0.291 |
| 4–5 | 0.067 | 1.17 | 1.19 | 0.276 | 1.000 |
| 4–6 | 0.338 | 2.18 | 6.68 | 0.010 | 0.146 |
| 5–6 | 0.271 | 1.87 | 4.13 | 0.042 | 0.505 |

**Table 2.** Pairwise comparisons among ecological guilds for residual running speed under the log-linear allometric PGLS model. Contrasts are from likelihood-ratio tests comparing guild-offset models, with Holm correction applied across all pairwise guild comparisons. The log_10_ contrast is defined as guild A minus guild B; speed ratios are therefore 10^Δ^. No pairwise contrast remained significant after Holm correction.

| Guild comparison | Δ log<sub>10</sub>(*v*) | Speed ratio | LR | *p* | *p*<sub>Holm</sub> |
|---|---|---|---|---|---|
| 1–2 | -0.170 | 0.68 | 1.68 | 0.195 | 1.000 |
| 1–3 | -0.274 | 0.53 | 3.84 | 0.050 | 0.554 |
| 1–4 | -0.271 | 0.54 | 3.98 | 0.046 | 0.554 |
| 1–5 | -0.211 | 0.61 | 2.66 | 0.103 | 0.907 |
| 1–6 | 0.081 | 1.20 | 0.20 | 0.656 | 1.000 |
| 2–3 | -0.103 | 0.79 | 2.69 | 0.101 | 0.907 |
| 2–4 | -0.101 | 0.79 | 2.61 | 0.106 | 0.907 |
| 2–5 | -0.041 | 0.91 | 0.53 | 0.465 | 1.000 |
| 2–6 | 0.251 | 1.78 | 3.50 | 0.061 | 0.614 |
| 3–4 | 0.002 | 1.01 | 0.00 | 0.976 | 1.000 |
| 3–5 | 0.062 | 1.15 | 0.73 | 0.391 | 1.000 |
| 3–6 | 0.354 | 2.26 | 6.14 | 0.013 | 0.185 |
| 4–5 | 0.060 | 1.15 | 0.94 | 0.331 | 1.000 |
| 4–6 | 0.352 | 2.25 | 7.35 | 0.007 | 0.101 |
| 5–6 | 0.292 | 1.96 | 4.85 | 0.028 | 0.360 |

